# Synthesis of titanium dioxide thin films and its application in reducing microbial load of milk

**DOI:** 10.1101/045179

**Authors:** Shruthi N Kumar, Manjula Sarode, Shankar H. N. Ravi

## Abstract

Milk has rich nutritional content packed with fats, proteins and water. It also contains beneficiary and non-beneficiary microbes which account for its short shelf life. Currently cold storage, pasteurization, ultra-high temperature, microfiltration and addition of lactose peroxidase are the methods of choice to control milk spoilage and prolong its shelf life. Their limitations include high energy consumption and loss of variable proportion of heat sensitive nutrients. Titanium dioxide (TiO_2_) nanoparticle coated thin film was used in food packaging industry for its antimicrobial property. TiO_2_ thin films were synthesized by sol gel process; it was characterized with scanning electron microscopy which showed pore size as 5 μm and Fourier transform infrared spectroscopy which showed metal present in sample is TiO_2_, zinc and silver. Exposure of raw milk at room temperature to TiO_2_ thin films doped with zinc or copper for a couple of hours showed zone of inhibition in disc diffusion technique, reduction in acid production. It also showed reduction in optical density indicating inhibition of growth in growth curve analysis, increase in the time required for methylene blue reduction and a five log folds decrease in bacterial count estimated using serial dilution plate count. The present studies were carried out under room temperature and pressure which is an added advantage in terms of energy as well as retention of nutrients. Though TiO_2_ is insoluble in water one needs to address it toxicity issues and adverse effects if any on the nutritional quality of milk before scaling up the process.

## 1 Introduction

Milk is a rich source of nutrients that helps the growth and maintenance of health of young ones and adults. The National Institute of Nutrition has recommended a minimum of 300 g daily intake of milk for children between 1-3 years of age and 250 g for those between 10-12 years. It contains 87% water, 4% fat and 9% solid non-fat which include proteins such as casein and albumin, and carbohydrates such as lactose to an extent of 5 % and around 0.7 % ash (Rhea Fernandes, 2009).

Apart from above mentioned nutrients, milk also consists many micronutrients which are essential for growth of children and adults. Milk contains nine essential nutrients like calcium, protein, vitamin A, vitamin B12, vitamin D, potassium, phosphorus, niacin, riboflavin. Spoilage bacteria may originate on the farm from the environment or milking equipment or in processing plants from equipment, employees, or the air. Microorganism effects quality of milk both physically and chemically in following ways:

**Table 1.**
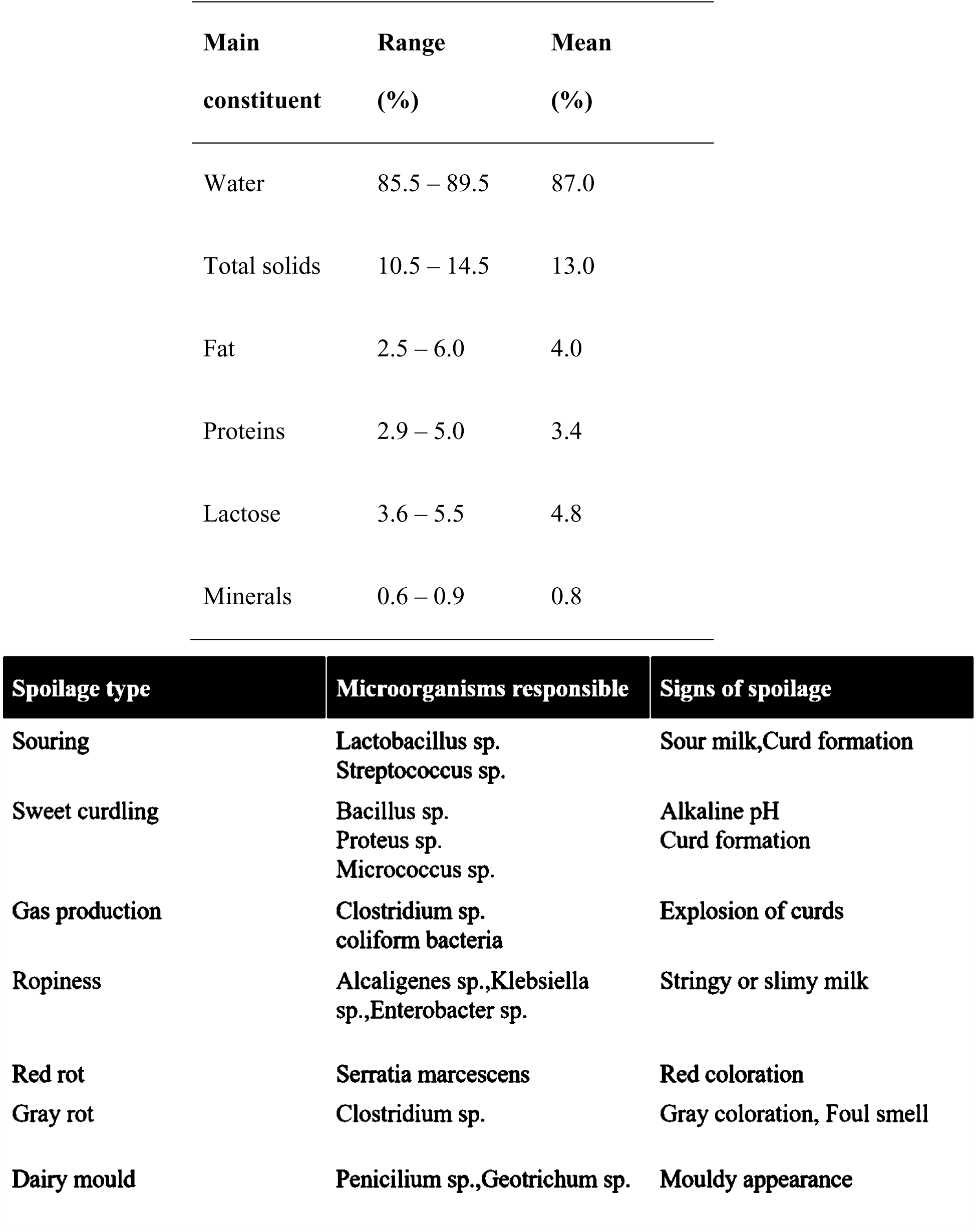
Table 1a: Cow milk composition (Robert G, 1995) Table 1b: Spoilage of microorganisms.

**Fig 1.**
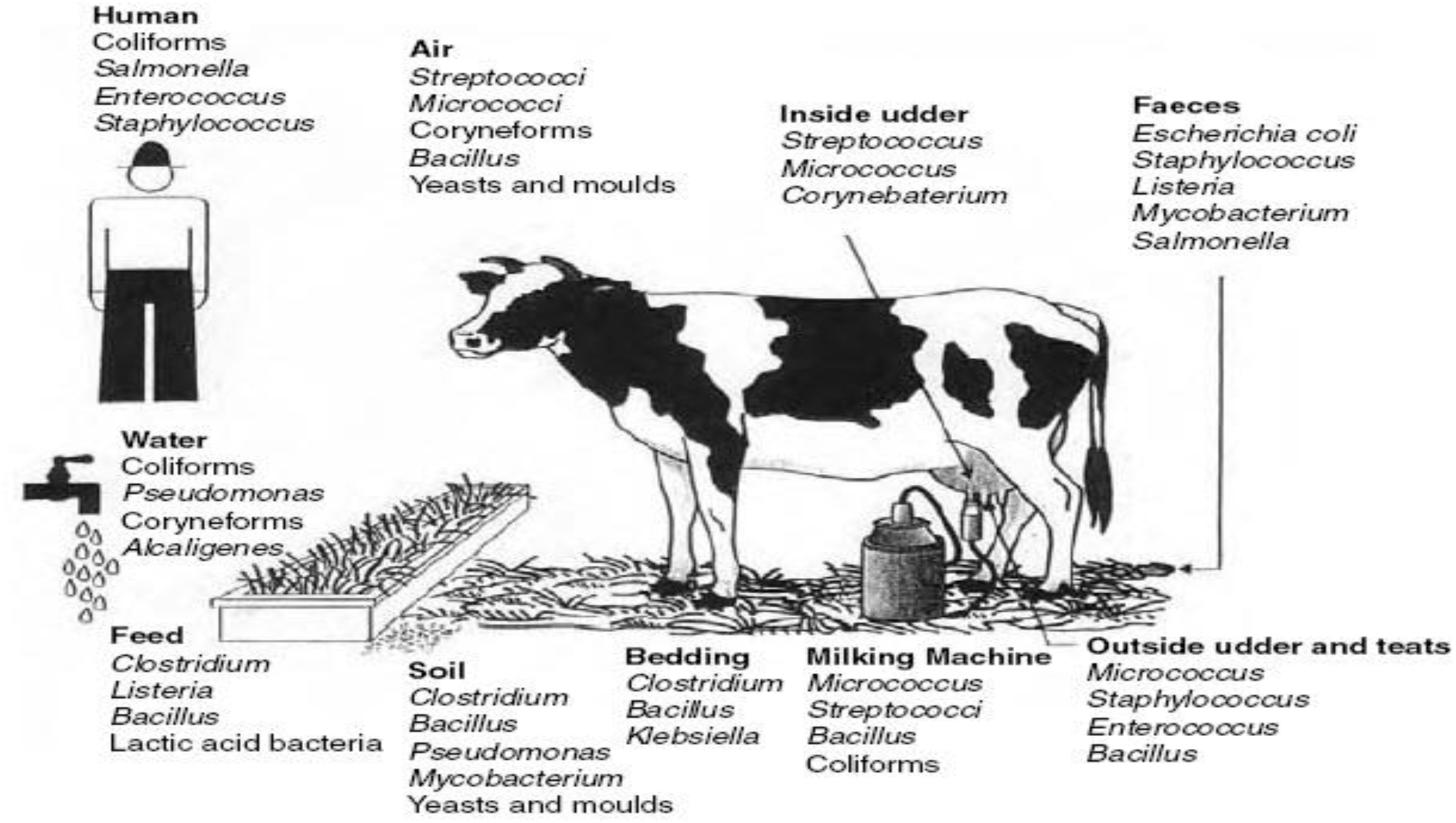
Sources of microbial contamination in milk (Loralyn H. Ledenbach and Robert T. Marshall, 2009)

### 1.1 Indian dairy industry

India is leading milk producer in the world and has about 15% of world’s livestock population. Milk production in India is greater than 16% (135 MT) of world’s production.

Indian dairy sector contributes majority share in domestic products. Presently India has about 69,000 dairy cooperatives which gives employment to more than 72 mn dairy farmers. Despite the increase in milk production in last three decades (reference: Operation Flood), milk yield per animal is still low. The main reasons for the low yield are lack of use of modern practices in milking, fodder in all the seasons, access to veterinary health services. Indian dairy has improved over years from importer to exporter that is per capita availability from 132 g/day to 302 g/day. Milk undergoes various processes before reaching the consumer and also there is a huge loss of produced milk (4).

**Table 1_1:** Main problems in Indian dairy industry is at various levels:

|  |  |
| --- | --- |
| Small holder level | At this level there is lack of good quality feed, veterniary facilities, chilling facilities. |
| Collection level | Here there is lack of infrastucture and also there are too many intermediates. Also there is manipulation of quality of milk by farmers |
| Processing level | Here there is seasonality of production and fluctuating supply, absence of quality standards, adulteration is more and lack of trained and skilled workers. |
| Storage and logistics level | Here lack of cold storage facilities and also gap in cold chain and transport facilities. |
| Co-operative level | Ther is lesser participation in the decision making process, inefficient services and lastly insufficient infrastructure. |
| Marketing level | There is lack of transparent milk pricing system. |

**Fig 1.1a.**
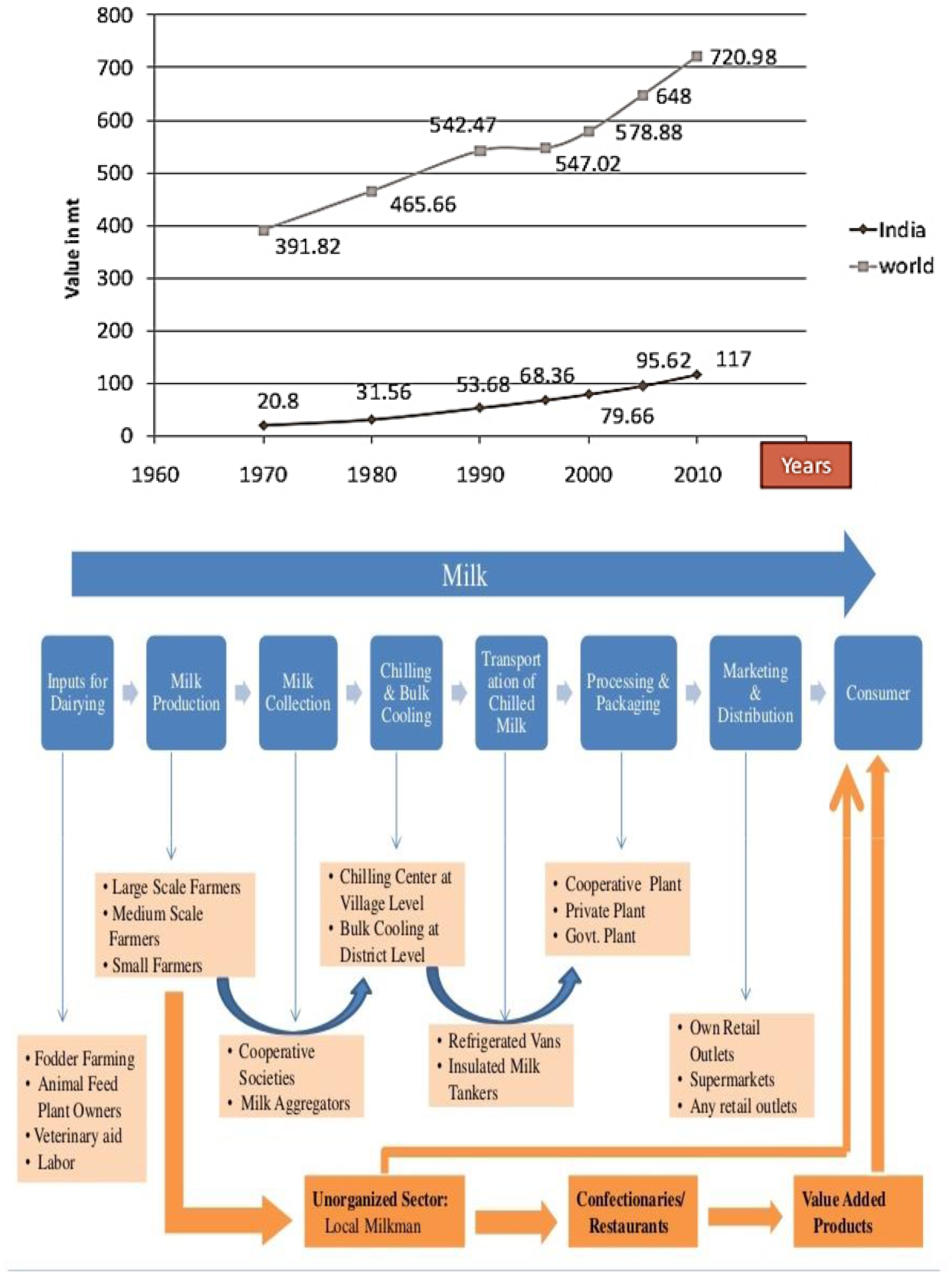
Trend in milk production world-India (Source-NDDB) Figure 1.1b: Supply chain in Indian dairy industry.

### 1.2 Milk processing

Milk is collected from farms and stored under chilled conditions at around 4°C, where mostly Pseudomonas species like P.fluorescens, P.fragi, P.lundensis survive and also stable proteolytic and lipolytic enzyme survive even in Ultra-High Temperature (UHT) which cause further spoilage. To control psychotrophs, normally industries follow various steps like thermisation where milk is heated to 57-68°C for 15-20 s followed by rapid cooling at <6°C. This eliminates lot of psychotrophic bacteria but not vegetative pathogens, for example Listeria monocytogenes which survive under UHT conditions and grow under chilled conditions (Mackey B.M, & Bratchell N. 1989; Bradshaw et.al,1991; Knabel et.al, 1991; Farber et.al,1992; Linton et.al, 1990). Deep cooling is the process where milk is stored at temperature as low as 4°C to improve the shelf life of milk. In carbon dioxide addition method, carbon dioxide is added at concentration of 20-30 mM (Amigo L & Calvo M.M.,1996; Joseph et.al 2006) which inhibits microbes by lowering pH or displacing oxygen or inhibiting production of enzymes but major concerns were these help in growth of *Clostridium botulinium* (Glass et.al, 1999).

Other methods that are more commonly used to increase the shelf life of milk are pasteurisation and UHT. In pasteurization, milk is heated either at low temperature for long time (LSLT, 63-65°C for 30 min) or high temperature for short time (HTST, 71-72°C for 15s) where majority of pathogens are killed but not spores and in case of UHT, milk is heated to 120 °C for 30 min where majority of spores are destroyed.

Apart from above practised methods newer methods to increase shelf life of milk are being experimented like silver nanorods (Padmanaban et.al, 2013) have been used in dairy industry with promising results, use of casein micelles as nutraceuticals (Semo et.al, 2007), alpha lactalbumin nanotube (J.F. Graveland-Bikker, C.G. de Kruif, 2006) and TiO_2_ thin films for food packaging (Siti et.al 2014) are recent advances.

In case of India, alternative methods need to be thought of apart from conventional methods for chilling. Major reasons being chilling requires lot of energy and uniterrupted power supply. Next being cows are milked twice a day except 15 days before calf is born and 5 days after its birth and milk needs to be chilled within 4 h of drawing since India being tropical country, chances of spoilage is relatively higher. Others reasons for the need of alternative method is distance between village and processing unit is quiet far. So alternative was Rapid Milk Chiller, thermal battery operated chiller which is efficient if produce is about 500l.

### 1.3 TiO_2_ thin films

Silver is antimicrobial and has various application but it is expensive, so cheaper alternatives were sought. TiO_2_ has properties of being antimicrobial, stable, cheaper, hydrophilic, non-toxic etc. makes it an alternative choice for silver. TiO_2_ exists in three phases; brookite (orthorhombic), anatase (tetragonal) and rutile (tetragonal) A wide range of techniques have been used for synthesis of thin films such as Hydrothermal methods, Electron beam evaporation, magnetron sputtering, solvo thermal synthesis and sol–gel methods.Among these techniques the sol–gel method offers several advantages like homogeneous films, simpler processing, variation of thickness and area of the films (Chopra, 1986; West, 2003).

### 1.4 Antimicrobial activity of TiO_2_

TiO_2_ was used in air, water and surface disinfection, and also in inhibition of foodborne microorganisms. The microorganisms include foodborne microorganisms such as E.coli, Bacillus cereus, Staphylococcus aureus, Norovirus and several salmonella strains etc.

**Fig 1.4.**
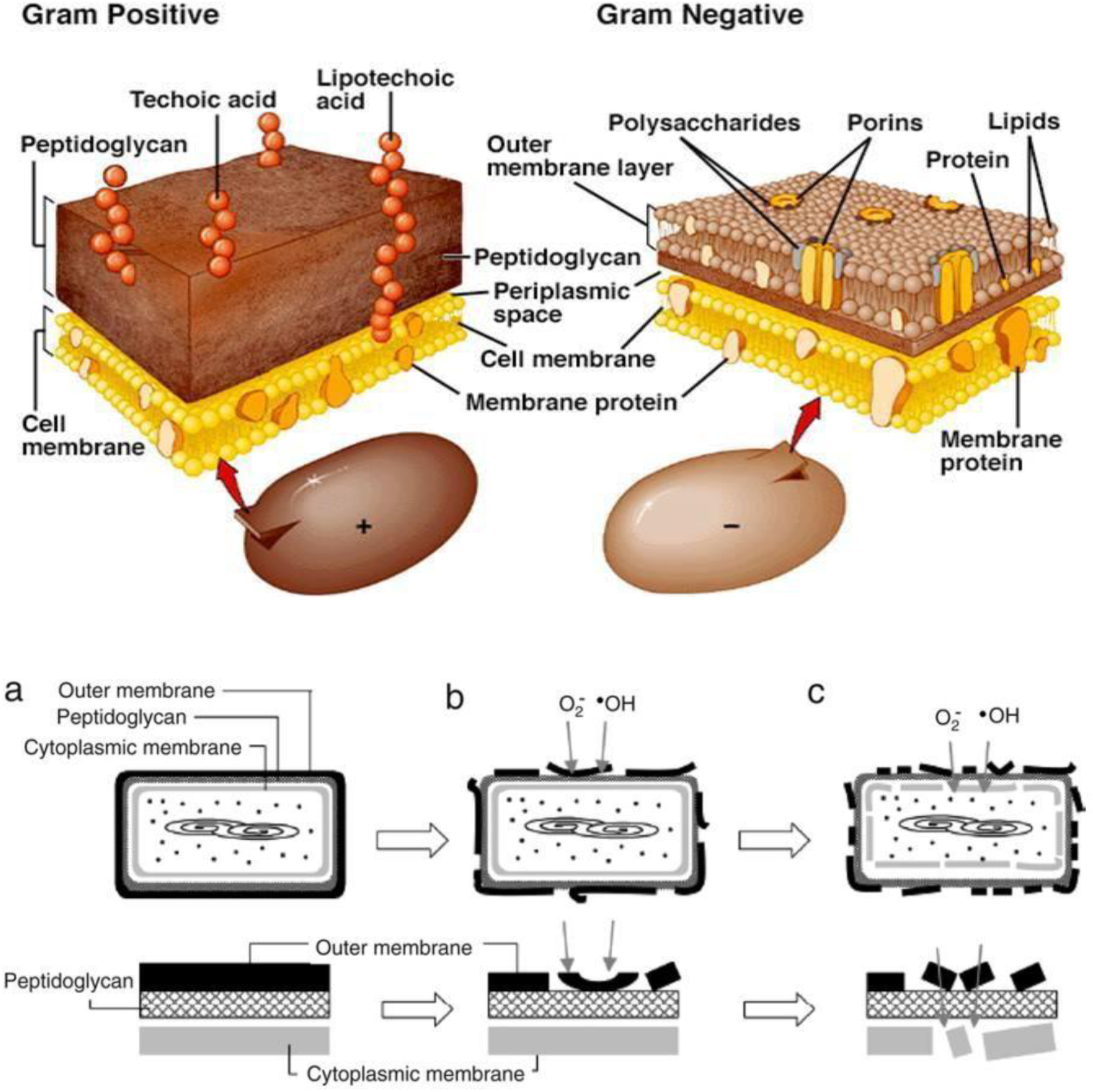
Fig 1.4 a - Cell wall constitution in gram-positive (left) and gram-negative (right) bacteria (adapted [21]). Fig 1.4 b - Schematic illustration of the stages in the process of E. coli photokilling on a TiO_2_ film. In the lower row, part of the cell envelope is magnified (adapted [21])

Gram-positive bacteria cell wall possesses a thick peptidoglycan layer (PG) and an inner membrane (cytoplasmic membrane or inner membrane, IM), while gram- negative cell wall is made of an outer membrane (OM), a thin PG layer and an IM. The differences in cell wall composition is reason for resistance to disinfection of gram-positive bacteria compared to gram-negative.

Another study of photokilling process of *E.coli* by using AFM showed disordering of the OM of bacteria cells by reactive species (•OH, H_2_O_2_, O_2_•–) (Figure 2.5 (ii)), which leads to changes in the permeability of ROS and, as a result, enable an easy attack to the inner membrane, resulting in the peroxidation of the lipids that constitute the cytoplasmic membrane. The destabilization of the cytoplasmic membrane leads to changes in the cell permeability, with the loss of ions (disruption of the electrochemical balance) and other important molecules, eventually resulting in loss of cell viability and ultimately cell death (Sunada et.al,2003).

### 1.5 Toxicology of TiO_2_

TiO_2_ is generally insoluble in water but TiO_2_ nanoparticles are expected to alter biological properties like bioavailability and cytotoxicity. Apart from National Institute for Occupational Safety and Health (NIOSH) recommended exposure limits (REL) no other exposure limits have been set and tolerance limits is 5-10 μg/ml. In certain invitro and invivo experiments, reproductive and developmental toxicity is studied but results are unclear. Accumulation of TiO_2_ in organs and tissues may take place on continuous exposure and also this may cause generation of reactive oxygen species (ROS) and alternate cell signal in pathways that play role in understanding behaviour of carcinogenesis. It also has shown to cross blood-brain, blood-placenta barriers. Toxicokinectic studies include absorption, distribution, excretion, accumulation and metabolism through different route in body. Further studies need to focus on systemic response from organ exposure and biomarker reflecting exposure and toxic effect (Hongbo Shi et.al,2013; Jinyuan Chen et.al,2009).

In the present work, TiO_2_ thin films were synthesised using sol gel process and doped with metals like zinc, copper. The synthesized films were used to study their effect on reducing microbial load of water and the studies were extended on milk samples.

## 2 Materials

Fresh cow milk was collected under hygienic conditions in clean vessels from a nearby farm and brought to the laboratory under cold conditions. Cow milk was allowed to rest under chilled conditions (4°C) for four hours before conducting the experiments in order to settle the gas produced during milking. Zinc nitrate, cuprous chloride, titanium isopropoxide, acetyl acetone, CTAB, silver nitrate, isopropyl alcohol, vanadium pentoxide, iodine, potassium iodide, potassium chromate, zinc chloride were of analytical grade and were obtained from Industrial labs, Bangalore. Nutrient broth, agar agar, methylene blue powder, phenolphthalein indicator, resorcinol was procured from Merck chemicals, Bangalore.

## 3 Methodology

### 3.1 Preparation of thin films

Titanium dioxide thin films was prepared using titanium isopropoxide solution and isopropanol in volume ratio of 1:5 in a beaker and the mixture was stirred for 10 minutes till yellow colour solution was obtained. Later 5ml of acetyl acetone solution was added to the above solution followed by addition of CTAB solution (0.03%) and the contents were stirred till clear solution was obtained (Isrihetty Senain et.al,2010).

### 3.2 Doping of Ti_2_ thin films

#### 3.2.1 Selection of dopants

Dopants were selected on basis of solubility in isopropanol solution and also their stability (24 hrs)(Ravishankar Rai V and Jamuna Bai A, 2011).

| Dopants | Result (solubility and stability) |
| --- | --- |
| Iron (Ferrous sulphate) | Negative (soluble, not stable) |
| Zinc (zinc nitrate) | Positive (soluble, stable) |
| Copper (cuprous sulphate) | Negative (partially soluble, not stable) |
| Nickel (nickel chloride) | Negative (not soluble, not stable) |
| Copper (cupric chloride) | Positive (soluble, stable) |
| Manganese (Manganese sulphate) | Negative (soluble, not stable) |
| Magnesium (Magnesium chloride) | Negative (not soluble, not stable) |
| Cobalt (cobalt oxide) | Negative (not soluble, not stable) |

Based on above criteria copper and zinc were chosen.

Metal dopants zinc and copper was added to thin films prepared in section 3.1 and the solution was used in synthesizing the films by coating on a microslide using spin coating apparatus (DELTA SPIN-1 RC2100XT543) at 300 rpm initially for few seconds, which was increased to 1000 rpm for 5 min. Then these films were dried in hot air oven at 60°C for 30 min and annealed at 200°C for 1 h in muffle furnace.

### 3.3 Testing for microbial and keeping quality of milk

#### 3.3.1 Disc diffusion test

Nutrient Agar was prepared using 2.72 g dehydrated nutrient broth powder in 200 ml of water mixed with 4 g (2%) of agar in 250 ml conical flask and autoclaved at 121°C, 15 PSI pressure for 15 min and on cooling was poured into sterile petri plates.*E.coli* culture suspension was spread on the surface of agar to develop a lawn of bacteria. Sterile Filter paper discs were loaded with 20 μl of test solution and placed on agar surface under aseptic conditions. Standard antibiotic disc was included as a positive control. The plates were incubated at 30°C for 48 h and observed for the presence of zone of inhibition.

#### 3.3.2 Growth curve analysis

Nutrient broth was prepared as in section 3.3.1 and was inoculated heavily with *E.coli* culture in 250 ml conical flask. The inoculum was incubated in an orbit shaker at 100 rpm at 30°C in presence or absence of titanium isopropoxide with and without dopants. The absorbance at 600 nm was read every 1 h in photoelectric colorimeter (Model: SYNTRONICS 112) with respect to control, blank and test compounds.

#### 3.3.3 Methylene blue reduction test

One ml of 1% methylene blue solution was added to 9 ml of water or milk to be tested in presence or absence of test compounds and the time taken for the disappearance of blue colour was recorded.

#### 3.3.4 Serial dilution Plate count

One ml of the sample to be counted is added to 9 ml of sterile 0.85% saline solution, which is vortexed and serially diluted in saline. An aliquot was spread in duplicate on nutrient agar plates and incubated at 30°C for 48 h. The number of bacterial colonies that developed were counted and used to calculate the bacterial numbers per ml of test sample.

**Fig 3.3.4.**
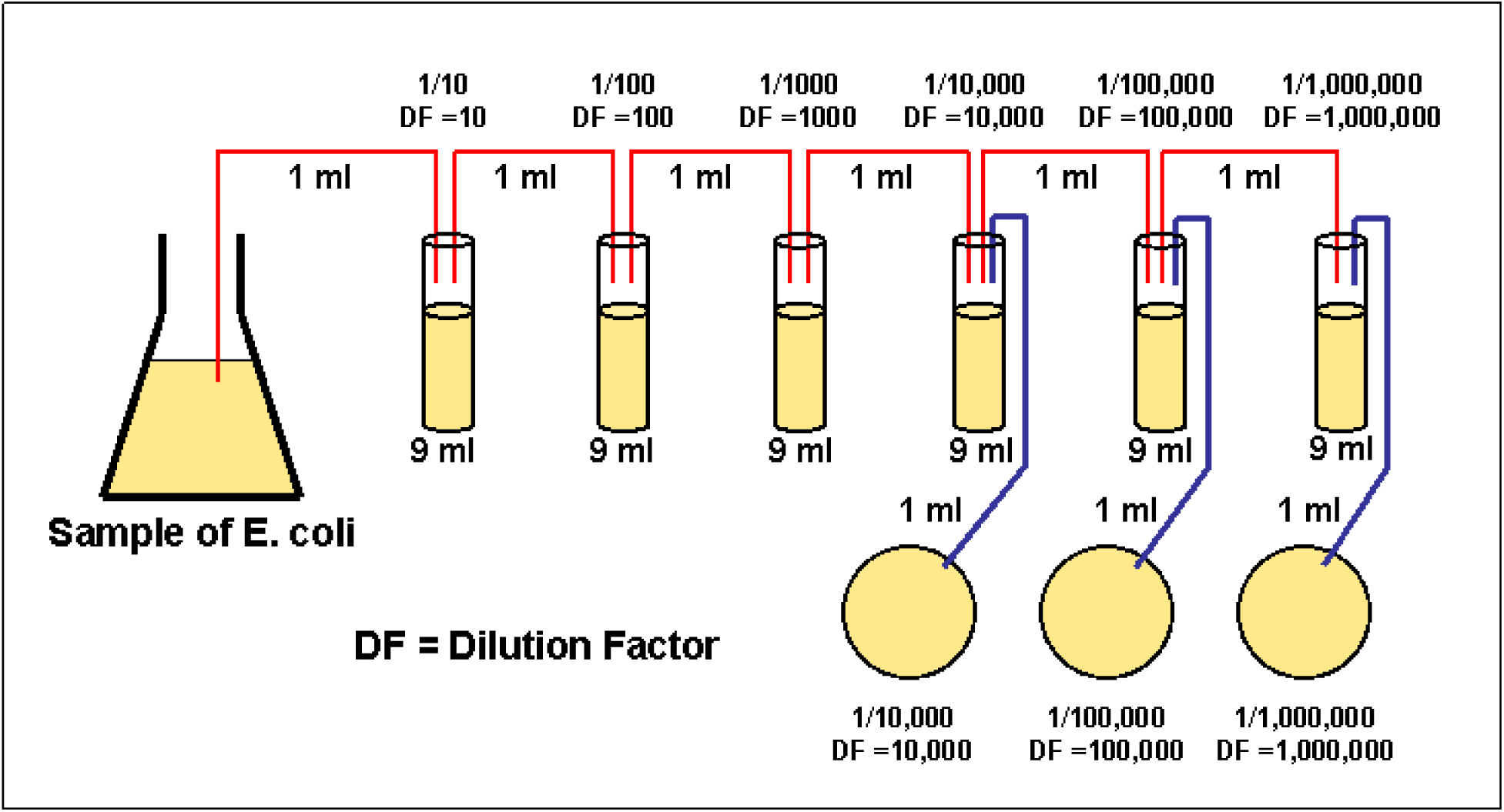
Serial dilution method

#### 3.3.5 Clot on boiling

Milk sample was taken in test tube and heated over flame in laminar airflow (LAF) to check whether milk curdles or not.

#### 3.3.6 Titrable acidity

An aliquot of milk sample was titrated against 0.01N NaOH with 2 drops of phenolphthalein as indicator and amount of NaOH consumed before the solution turned pink was noted and was calculated as:

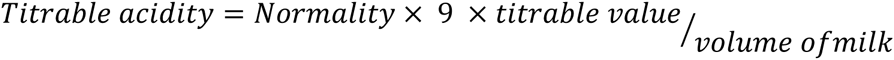

#### 3.3.7 Quality Analysis of Raw Milk Sample

a) Detection of Cane Sugar by using Modified Seliwanoff’s method: 0.5 g of resorcinol was dissolved in 40 ml of distilled water. To this 35 ml of conc HCl was added and make up volume to 100 ml using distilled water. 1ml of milk and 1ml of resorcinol solution was taken in test tube and placed in boiling water bath for 5 minutes and change in colour (either cherry red or pink) was observed.
b) Test for presence of Hydrogen Peroxide: 1 g of vanadium pentoxide was dissolved in 100 ml dilute sulphuric acid (6 ml concentrated sulphuric acid was diluted to 100 ml). 10-20 drops of reagent were added to 10 ml milk sample and mixed and colour change was observed that is red or pink.
c) Detection of Cellulose in Milk: Iodine-zinc chloride reagent was prepared by dissolving 10 g of zinc chloride in 4.25 ml water and after cooling iodine solution was added drop wise till it precipitates out. Iodine solution was prepared by mixing 1.5 g potassium iodide with 0.75 g of iodine in 30 ml water. 10 ml of milk was taken in a 100 ml beaker. To this 50 ml of hot water was added and stirred thoroughly for about 2 min until it dissolved. The mixture was filtered with filter paper twice using hot water. Residue was scraped and placed on a glass slab and colour change was observed that is blue colour in iodine-zinc chloride region and no blue colour in iodine region.
d) Detection of Sodium Chloride in milk: 5 ml of milk sample was taken in a beaker and 1 ml of 0.1 N silver nitrate solution was added to it. The contents were mixed thoroughly and 0.5 ml of 10% potassium chromate solution was added to it. The colour of resulting solution was observed; appearance of yellow coloured solution will indicate presence of chlorine.
e) Detection of Starch: Iodine solution was prepared by dissolving 1.3g of iodine and 1.5g of potassium iodide and volume was made up to 100 ml using distilled water. About 5 ml of milk sample was taken in a test tube. Milk was boiled and test tube was allowed to cool to room temperature. 1-2 drops of iodine solution were added to the test tube. Development of blue colour, which disappears when sample is boiled and reappears on cooling, indicates presence of starch.
f) Total Solid Fat content: The weight of empty crucible was noted.5ml of milk sample was taken in it and weighed (W1). It was placed in hot air oven at 110° C. After one hour the sample was taken out and weighed again (W2). Total solids were determined as a percentage of the difference between these two weights.
g) Specific Gravity of samples: The raw milk sample was filled in a specific gravity bottle. The weight of the bottle with and without sample was noted.
h) pH of samples: The sample was poured into a sterile beaker and placed in a water bath at room temperature, its pH was measured using pH meter.

## 4 RESULTS

### 4.1 Characterisation results

Thin films of TiO_2_ doped with zinc and copper were prepared as described in methods section 3.1 and 3.2 and characterized using FTIR and SEM techniques.

#### 4.1.1 FTIR

Fig 4.1.1 shows the Infra red spectral scan of TiO_2_ doped with zinc. Two peaks were observed at 700 and 1000 cm^-1^. This means it contains functional groups of OH and CH3 obtained from isopropanol.Figure 4.1.2 shows the spectral scan of TiO_2_ doped with zinc and silver. Peak broadening may be observed at 1000 to 1220 cm^-1^. This is possibly due to presence of zinc.

**Fig 4_1_1.**
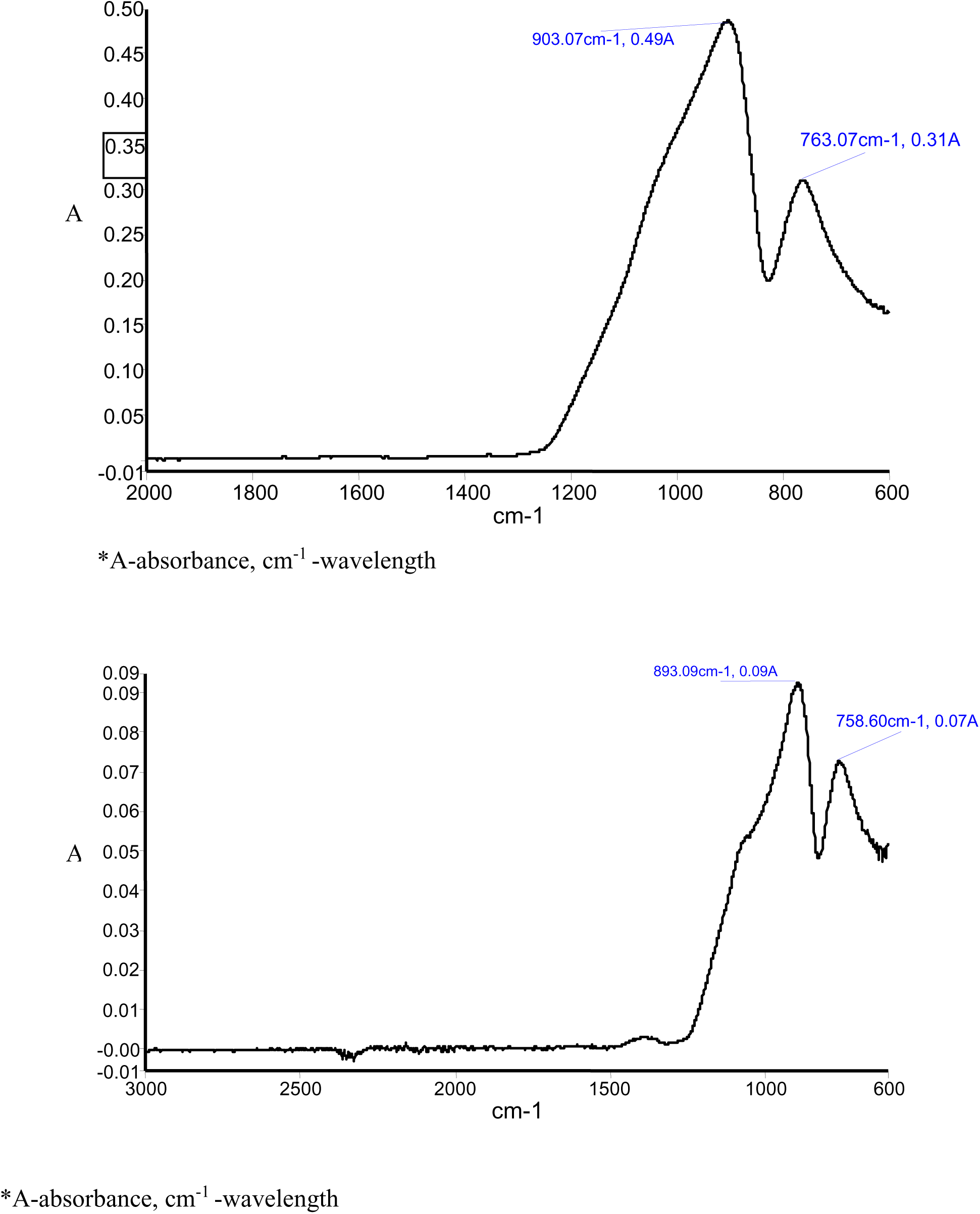
Fig 4.1.1a: FTIR spectral scan of TiO2 doped with zinc Fig 4.1.1b: FTIR spectral scan of TiO_2_ doped with zinc and silver These tests show the presence of TiO_2_, Zn, Ag.

**Fig 4.1.2:**
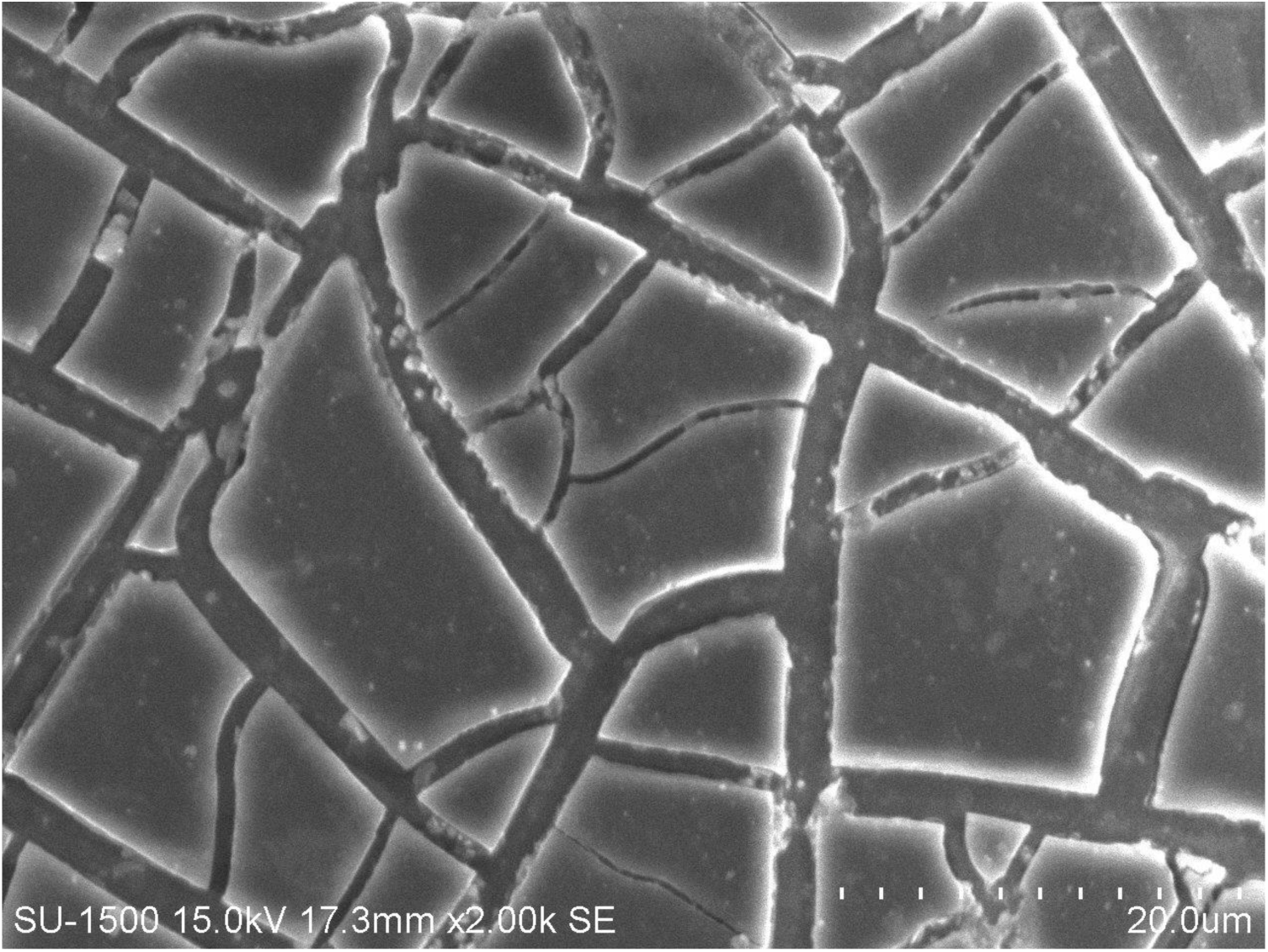
SEM image of TiO_2_ doped with zinc and silver.

**Fig 4_2_1.**
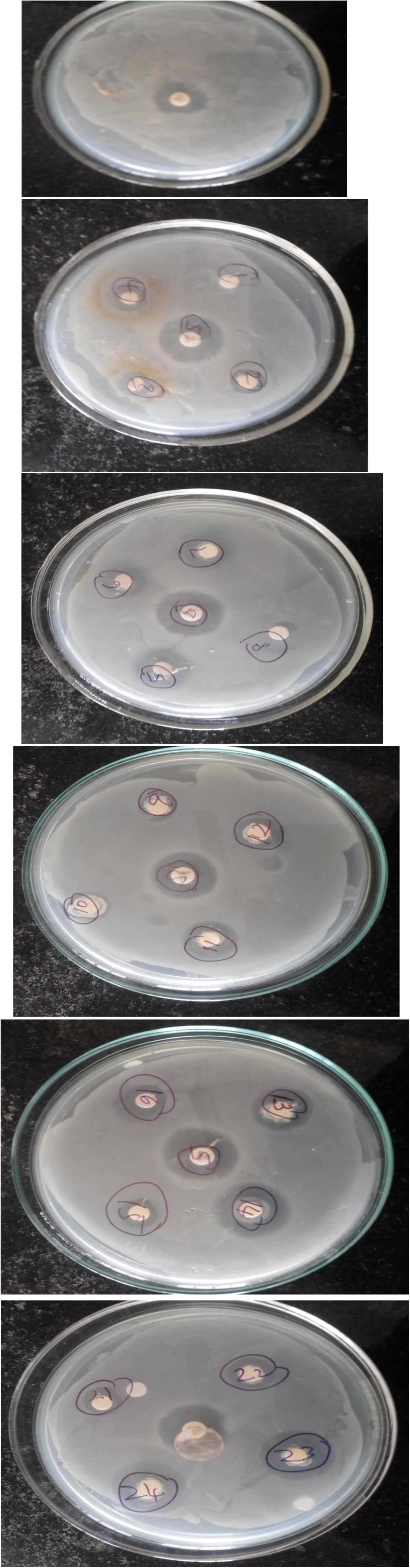
Fig 4.2.1a: Standard disc (Ampicillin 10 mg as positive control). Fig 4.2.1b: Treatment 1: Isopropanol; Treatment 2:0.02g of copper/ml of titanium isopropoxide; Treatment 3:0.04g of copper/ml of titanium isopropoxide; Treatment 4:0.06g of copper/ml of titanium isopropoxide; standard. Fig 4.2.1c: Treatment 5:0.08g of copper/ml of titanium isopropoxide; Treatment 6:0.1g of copper/ml of titanium isopropoxide; Treatment 7: Titanium isopropoxide; Treatment 8: isopropanol; standard. Fig 4.2.1d: Treatment 9:0.02g of zinc/ml of titanium isopropoxide; Treatment 10:0.04g of zinc/ml of titanium isopropoxide; Treatment 11:0.06g of zinc/ml of titanium isopropoxide; Treatment 12:0.08g of zinc/ml of titanium isopropoxide; standard. Fig 4.2.1e: Treatment 17:0.06g of copper/ml of titanium isopropoxide; Treatment 18:0.08g of copper /ml of titanium isopropoxide; Treatment 19:0.1g of copper/ml of titanium isopropoxide; Treatment 20:0.02g of zinc/ml of titanium isopropoxide; standard. Fig 4.2.1f: Treatment 21:0.04g of zinc/ml of titanium isopropoxide; Treatment 22:0.06g of zinc/ml of titanium isopropoxide; Treatment 23:0.08g of zinc/ml of titanium isopropoxide; Treatment 24:0.1g of zinc/ml of titanium isopropoxide; standard.

#### 4.1.2 SEM

SEM image is taken at magnification of 500x resolution and it was also observed that there is lot of crack on thin film which could be due to lot of heat generated on impact of electrons with the sample. Size of microbes normally varies from 0.5μm to 2μm whereas approximate pore size analysed based on SEM image is 0.5μm

### 4.2 Effect of exposure Ti_2_ thin films on microbial load of water

Drinking water is normally clear and free of contamination but in some cases it can also contain heavy metals, microbes, etc. As water acts as carrier media for microbes, the effect of the synthesized thin films on water was studied to reduce the microbial load reduction.

The change in microbial load of water samples after its exposure to TiO_2_ thin films was assessed by disc diffusion test, growth curve analysis, methylene blue reduction test and serial dilution plate count.

#### DISC DIFFUSION TEST

*E.coli* was used as a test organism, to study the effect of various concentrations of zinc and copper as dopants in titanium isopropoxide solution which was tested by disc diffusion test. When sterile filter paper disc impregnated with a known concentration of test solution is placed on agar plate, immediately water is absorbed into the disc from the agar.

The result of disc diffusion test is shown below. It appears that titanium isopropoxide by itself inhibits the growth of *E.coli* and addition of zinc and copper at 2%, 4%, 6%, 8% and 10% improve the degree of inhibition since there was increase in the diameter of zone of inhibition.

Titanium isopropoxide showed inhibition of growth of *E.coli* which was accentuated by addition of zinc and copper as dopants. Isopropanol showed no zone of inhibition; This test is done in order to optimise concentration of dopant which was 10% in case of zinc and 8% in case of copper. Zinc at 10% was chosen for further testing as it showed better zone of inhibition than copper. This was studied in comparison with standard antibiotic disc here ampilicin was used.

#### 4.2.1 Growth curve analysis

The effect of exposure of water to titanium isopropoxide with and without doping was studied using growth curve analysis. Figure 4.2.2 shows the growth curve with respect to time. The result clearly demonstrates the inhibition of growth of *E.coli* in all the treatments. It shows titanium isopropoxide doped with zinc thin films showed better inhibition when compared to titanium isopropoxide or titanium isopropoxide doped with copper. Inhibition can be easily distinguished with control where bacteria were allowed to grow in media.

**Fig 4.2.2.:**
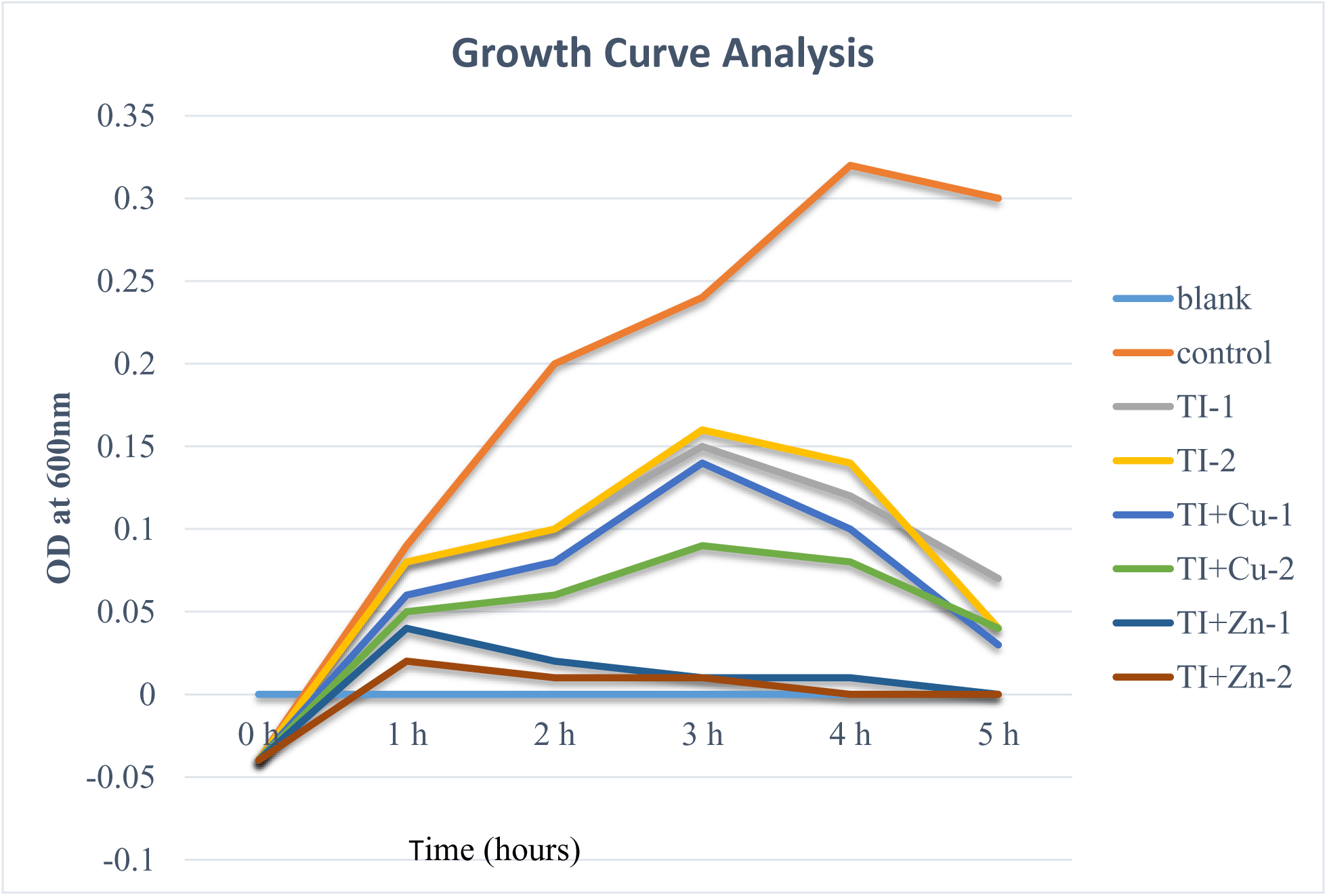
Growth curve analysis of *E.coli* as affected by zinc and copper doped TiO_2_

This test was done to in order to know minimum time required for inhibition when compared to control that is 2 h. Growth curve analysis of *E.coli* grown in broth in absence or presence of titanium isopropoxide with and without dopants showed that titanium isopropoxide reduced the bacterial growth. Addition of dopants increased the degree of inhibition. Zinc reduced bacterial growth in two hours when compared to copper and titanium isopropoxide which took longer time to inhibit. Control grew for some time and then curve began to drop which showed inhibition due to lack of nutrients Zinc is proven to be toxic to most of pathogens when compared to that of copper which inhibit only a few strains of bacteria.

#### 4.2.3 Methylene blue

This is preliminary test done in order to know quality of milk and if it is fit for consumption or not. The effect of addition of doped TiO_2_ on methylene blue reduction test was carried out and the results showed that there is a disappearance of blue colour in 2 h for raw water (inoculated with *E.coli* culture) while TiO_2_ treated water took 7 h to achieve same level of reduction (Fig 4.2.3).

**Fig 4.2.3:**
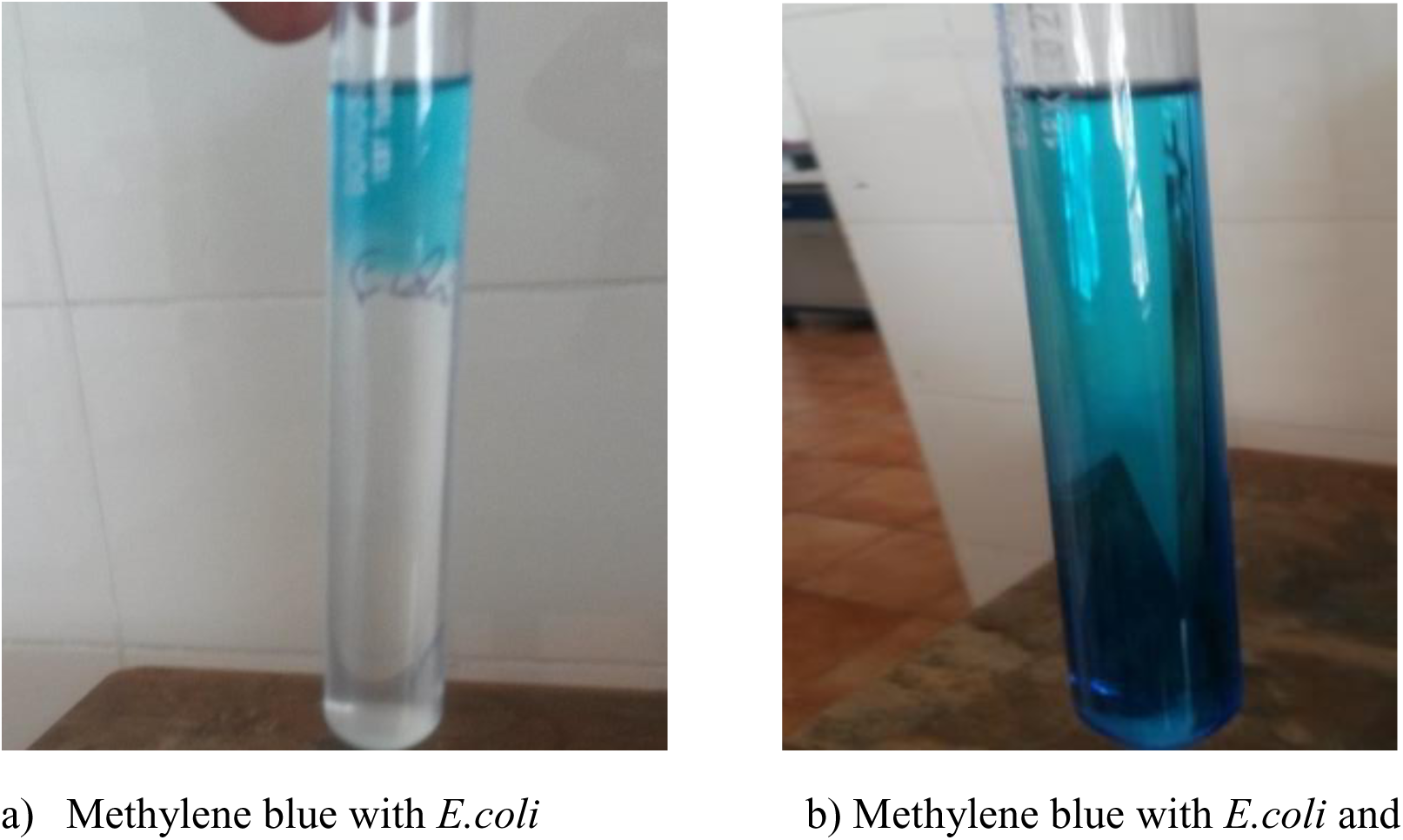
Methylene blue reduction test of water: a) Methylene blue with *E.coli* and b) Methylene blue with *E.coli* and TiO_2_.

**Fig 4.2.4:**
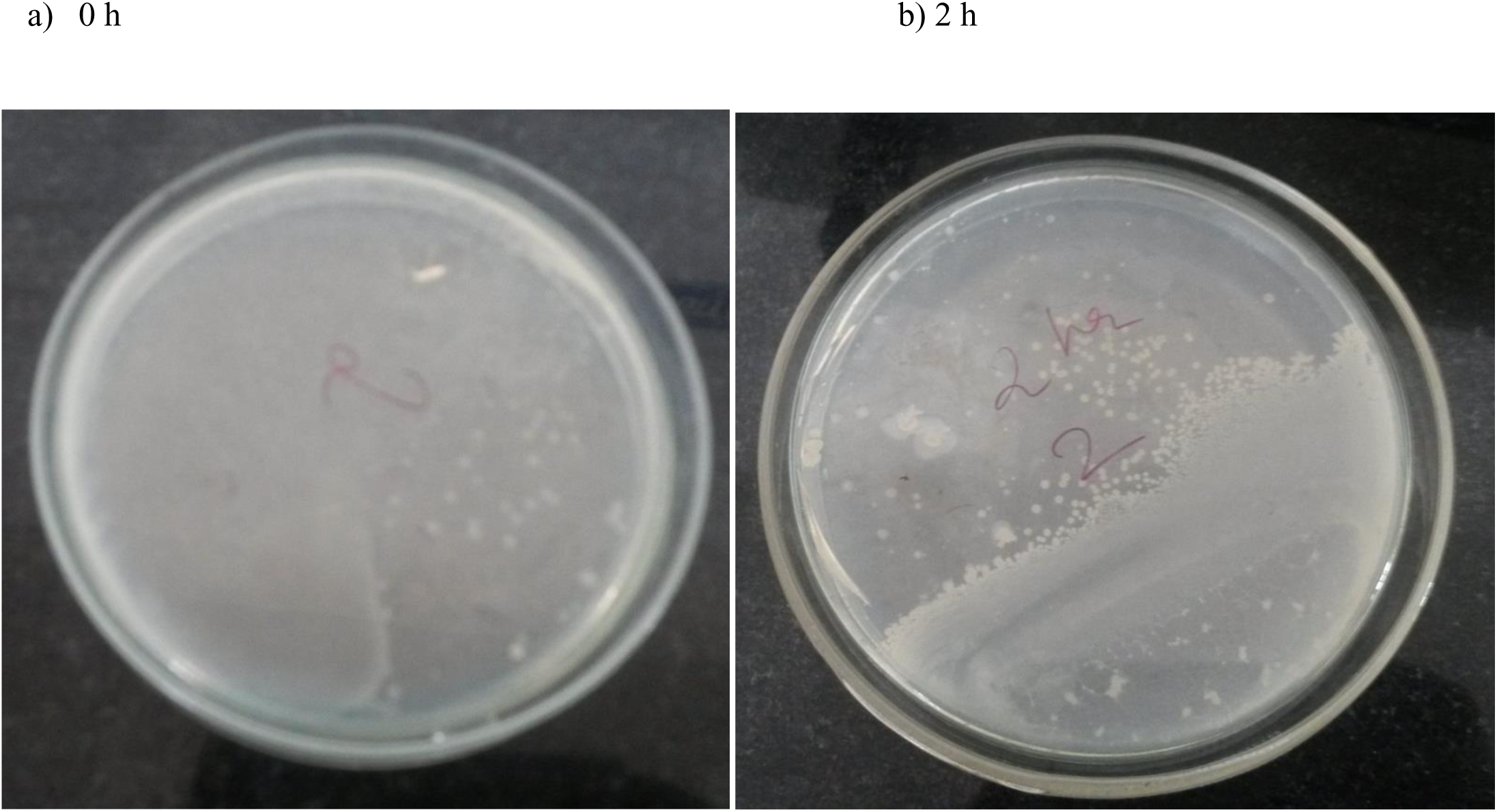
plate count a) 0 h b) 2 h)

Methylene blue with *E.coli* showed colour change from blue to colourless showing that*E.coli* activity reduces methylene blue within 2 h. Blue colour reappears on the top interface due to reoxidation of methylene blue. Methylene blue with *E.coli* and TiO_2_ showed no colour change as there was inhibition of *E.coli* even after 24 h. Water contaminated with culture could reduce methylene blue in 2 h suggesting high bacterial activity. Addition of zinc doped TiO_2_ increased the time of reduction to 7 h indicating loss of microbial number / activity. Addition of doped TiO_2_ alone to methylene blue without culture in beaker did not cause reduction supporting the idea that the reduction is not chemical but microbial in nature.

#### 4.2.4 Serial dilution method

This test is used to determine number of colonies present in given sample. Thin film of TiO_2_ coated on 7.5 cm x 2.5 cm microscopic slide was dipped into 200 ml water for 30 min.Table 4.2.4 shows the results of microbial counts. It can be observed that nearly five log fold reduction in bacterial population has occurred within a short exposure time (2 h). The result of exposure to TiO_2_ thin film on bacterial numbers as estimated by serial dilution plate count is shown in table 4.2.4.

**Table 4.2.4:**
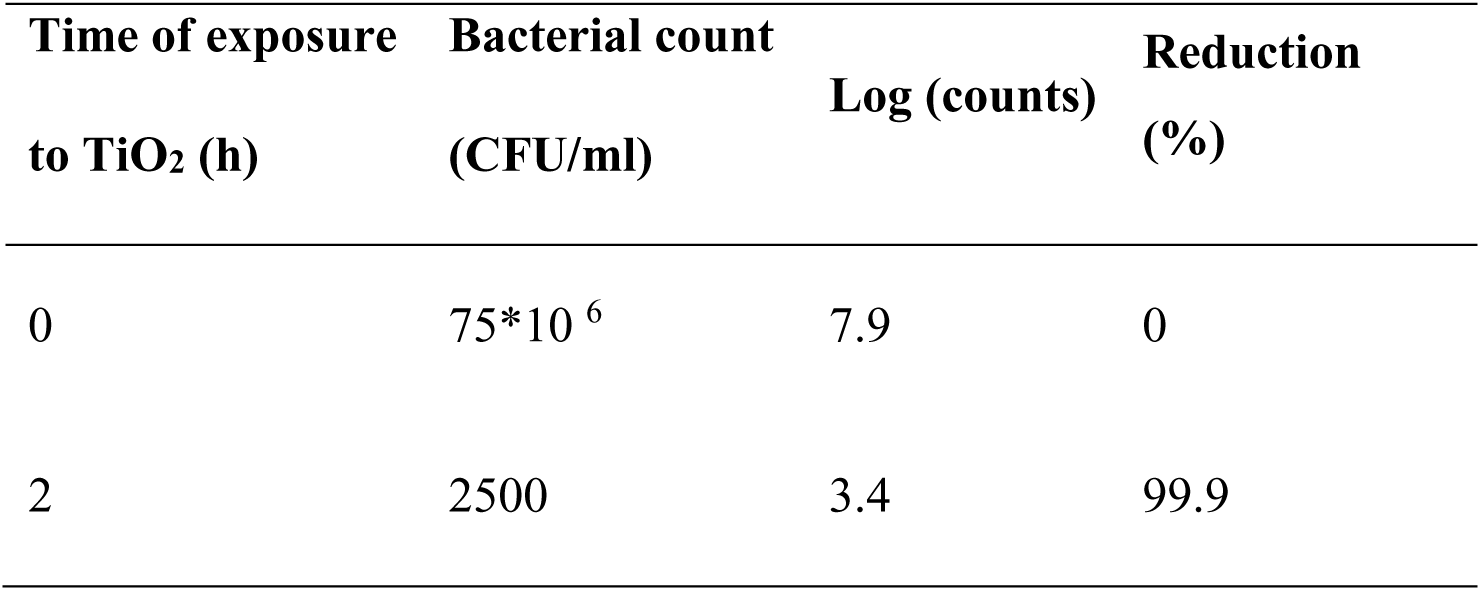
Effect of TiO_2_ thin film on microbial population of tap water for 30 min measured by serial dilution plate count.

### 4.3 Effect of doped Ti_2_ thin films on microbial load of milk

The effect of zinc and zinc with silver doped TiO_2_ thin films on the quality of milk was tested using clot on boiling, titrable acidity, methylene blue reduction and plate count. Consolidated results are shown in Table 4.3. Control milk curdled which was result of high titrable acidity value and also methylene blue reduced in 2 h attributed to microbial activity. Exposure of milk to zinc and zinc with silver doped TiO_2_ thin film showed low titrable acidity value which was insufficient to curdle milk and also methylene blue reduced in 4-8 h indicating loss of microbial activity. This is also supported by decrease in bacterial counts from 8 million at zero time to 2-2000 in 2 h.

**Fig 4.3.1:**
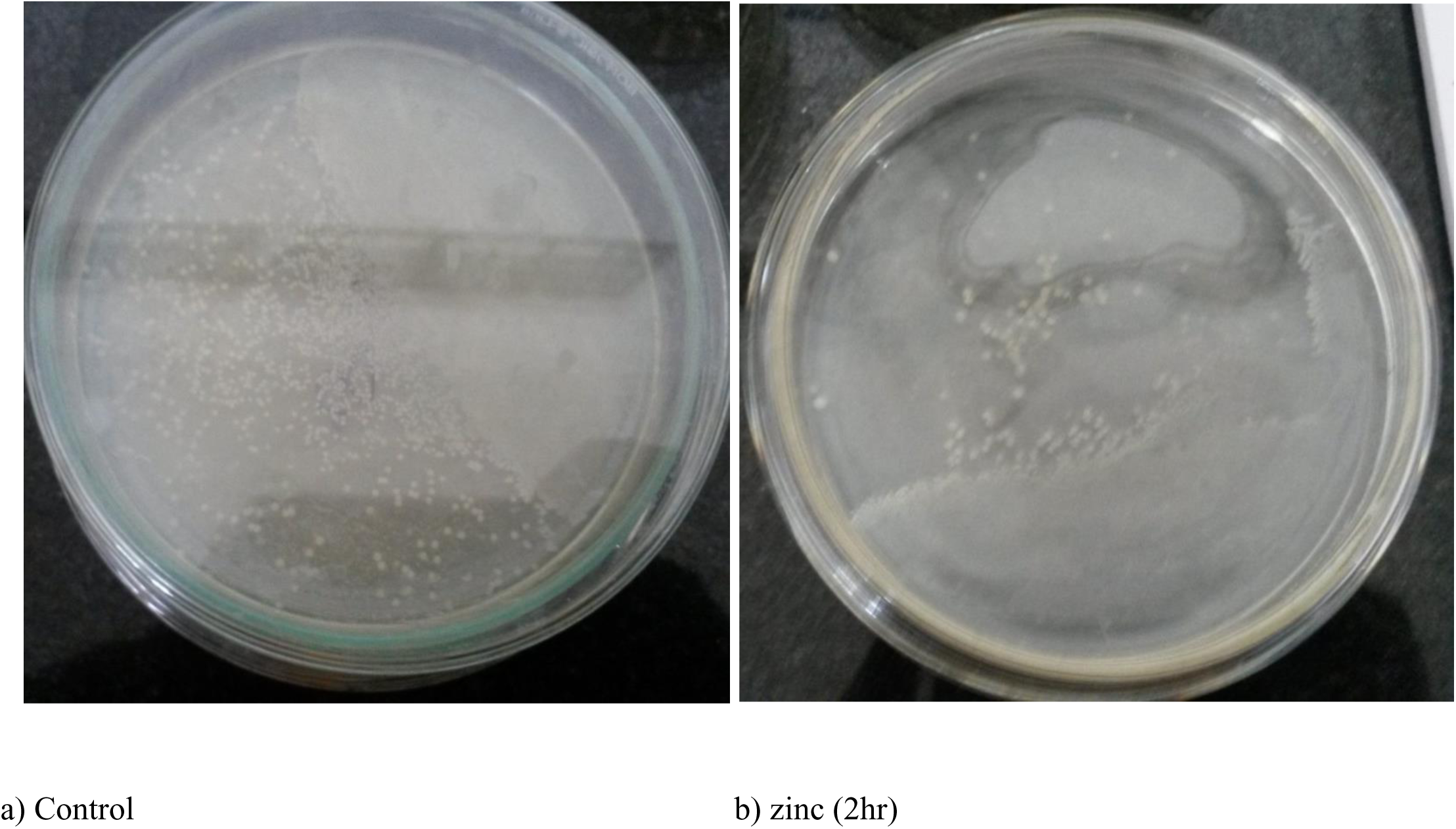
Colony count a) control b) zinc (2hr)

**Table 4.3:** Effect of zinc and zinc with silver doped TiO_2_ thin films on quality of milk.

|  | Control | Zinc (1 h) | Zinc with silver<br>(1 h) | Zinc (2 h) | Zinc with silver<br>(2 h) |
| --- | --- | --- | --- | --- | --- |
| COB | Curdling | No ppt | No ppt | No ppt | No ppt |
| Acidity<br>(percentage) | 0.225 | 0.135 | 0.09 | 0.079 | 0.045 |
| MBR (h) | 2 | 6 | 8 | 7 | 4 |
| Count (CFU/ml) | 80,00,000 | 3,50,000 | 20 | 2,000 | 2 |

**Fig 4.3.2:**
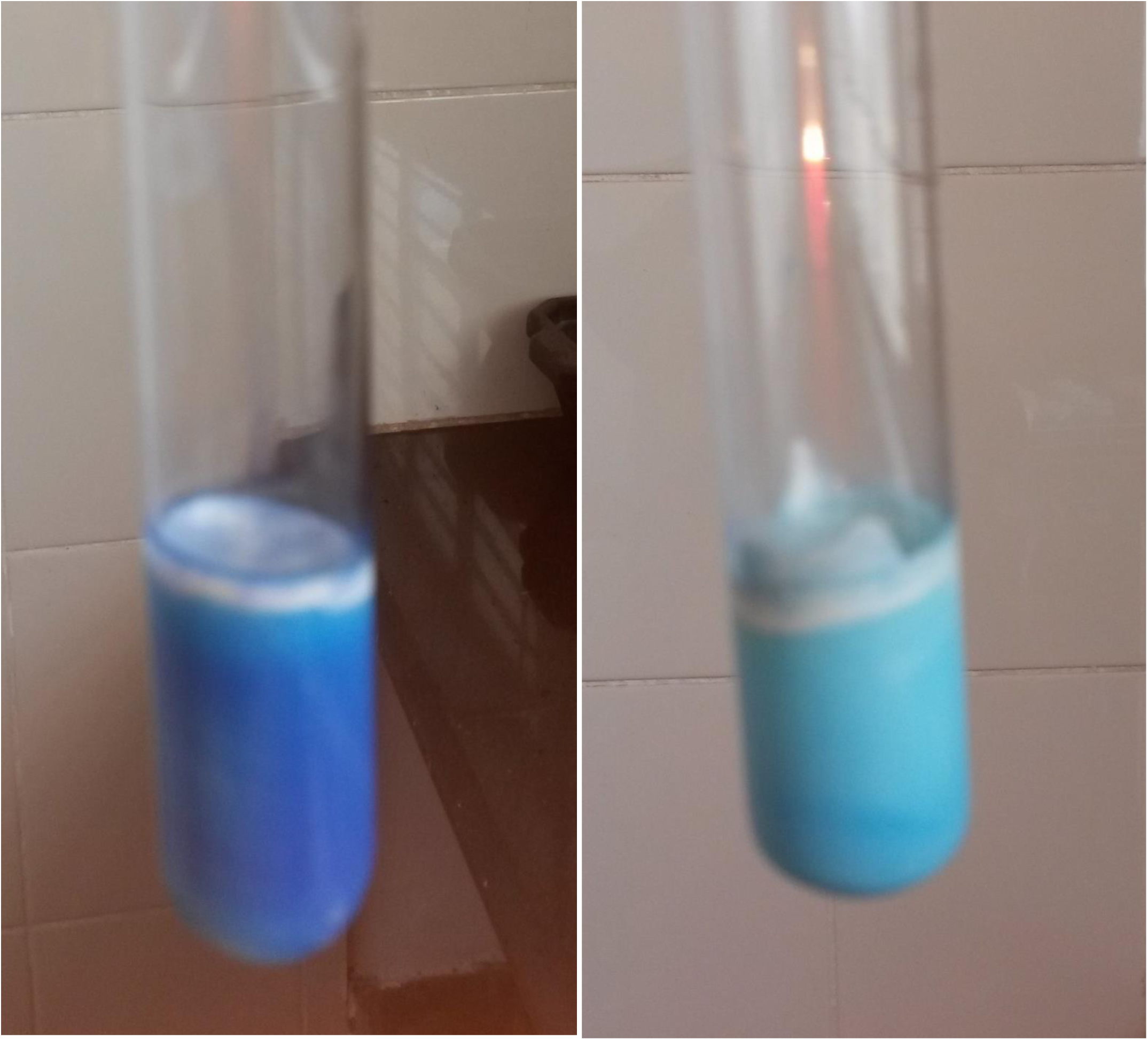
Methylene blue

The effect of doped TiO_2_ thin films on quality of milk was tested. Control milk reduced methylene blue in 2 h indicating high microbial load. Addition of doped TiO_2_ increased the time for reduction of methylene blue which indicated that there was a reduction in microbial number / activity.

Milk is a good medium for the growth of microorganism. A variety of microorganism can be found in both raw milk and pasteurized milk. These microbes reduce the oxidation reduction potential of the milk medium due to the exhausted oxygen by the microorganism. Milk is contaminated during the milking process, use of unsterilized dairy utensils such as milking machines, milk cans etc. Methylene blue reductase test is used to determine the quality of the milk. The principle of methylene blue reduction test is that the colour imparted to the milk by adding a dye like methylene blue will disappear more or less quickly, which depends on the quality of the milk sample to be examined. Methylene blue is a redox indicator that lose its colour under the absence of oxygen. The depletion of oxygen in the milk is due to the production of reducing substances in the milk due to increase in rate of bacterial metabolism. The dye reduction time refers to the microbial load in the milk and the total metabolic reactions of the microorganism. The presence of light fastens the reduction rate; hence the test tube under observation should be tightened properly.

Titrable acidity of milk has long been recognized and employed as an indicator of quality (Griffiths et al, 1988). It is expressed in terms of percentage lactic acid since lactic acid is the principal acid produced by fermentation after milk is drawn from the udder. Fresh milk, however, does not contain any appreciable amount of lactic acid and therefore an increase in acidity is a rough measure of its age and bacterial activity. Within a short time after milking, the acidity increases perceptibly due to bacterial activity. The degree of bacterial contamination and the temperature at which the milk is kept are the chief factors influencing acid formation. Therefore, the amount of acid depends on the cleanliness of production and the temperature at which milk is kept. For this reason, determination of acid in milk is an important factor in judging milk quality. Acidity affects taste as well. When it reaches about 0.3%, the sour taste of milk becomes sensible. At 0.4% acidity, milk is clearly sour, and at 0.6% it precipitates at normal temperature. At acidity over 0.9%, it moulds. Milk contains energy sources such as lactose (sugar), nitrogenous compounds such as proteins, amino acids, ammonia, urea etc. for the growth of microorganism.

Clot on boiling milk occurred after 2 h in control milk. While no such precipitation was observed in TiO_2_ treated milk. Control milk had highest titrable acidity while doped TiO_2_ treated samples showed variable amounts of acid; which is insufficient to curdle the milk. Bacterial population as measured by serial dilution plate count also decreased by five log folds with zinc doped TiO_2_ and while zinc with silver doped TiO_2_ showed drastic reduction to double digits. Silver is a known antibacterial substance and hence this result is not surprising.

### 4.4 Quality test in milk sample

Milk was treated with TiO2 thin films doped with zinc and were tested to check if these films had any effect on physico-chemical properties of milk sample (30).

#### 4.4.1 Detection of Cane Sugar by Modified Seliwanoff’s method (qualitative method)

Sucrose is absent in milk and its presence indicates adulteration. The acid hydrolysis of polysaccharides and oligosaccharides yields simpler sugars followed by furfural as by-product. The dehydrated ketose then reacts with the resorcinol to produce a deep cherry red colour. In case of aldoses, they react to produce a faint pink colour. When TiO_2_ treated milk sample was tested for sugar, there was no colour change indicating absence of sucrose. In pure milk sample no colour change is observed. (*Reference: - IS 1479 (Part I) 1961 (Reaffirmed 2003) Methods of test for Dairy Industry – Rapid Examination of Milk. Bureau of Indian Standards, New Delhi*).

#### 4.4.2 Test for presence of Hydrogen Peroxide

Hydrogen Peroxide is used in preserving milk and also as a bleaching agent. Presence of hydrogen peroxide would increase pH and antibacterial activity. When treated sample was checked there was no colour change indicating absence of hydrogen peroxide (*Reference: - A.O.A.C 17thedn, 2000 Official Method 957.08 Hydrogen Peroxide in milk*).

#### 4.4.3 Detection of Cellulose in Milk

Sample when treated with thin films showed no colour change indicating absence of cellulose (*Reference: - Manual Methods of Analysis for Adulterants & Contaminants in Foods. I.C.M.R 1990page 27*).

#### 4.4.4 Detection of Sodium Chloride in milk

Absence of sodium chloride was observed by no change in colour in treated sample. (*Reference: - Pearson’s Composition and Analysis of Foods, 9*^*th*^*edn,1991 - Modified Mohr method, page 14*).

#### 4.4.5 Detection of Starch

Test for starch was found negative as there was no colour change in treated sample. (*Reference: - IS 1479 (Part I) 1961 (Reaffirmed 2003) Methods of test for Dairy Industry - Rapid Examination of Milk. Bureau of Indian Standards, New Delhi*).

#### 4.4.6 pH of raw milk sample

The control and sample were kept under ambient room temperature of 26.5°C for a period of 14 hours and pH of samples were measured at regular intervals. pH was found to be 6.61 which was within the specified range. pH indicate that solution was stable over period of time and thin films had no effect on its acidity.

| Time (min) | pH of Control Sample | pH of Treated Sample |
| --- | --- | --- |
| 0 | 6.61 | 6.61 |
| 60 | 6.61 | 6.61 |
| 200 | 6.61 | 6.61 |
| 250 | 6.61 | 6.61 |
| 300 | 6.61 | 6.61 |
| 450 | 6.62 | 6.61 |
| 600 | 6.52 | 6.61 |
| 650 | 6.41 | 6.55 |
| 700 | 6.40 | 6.52 |
| 750 | 6.32 | 6.51 |
| 800 | 6.31 | 6.50 |
| 850 | 6.30 | 6.50 |
| 900 | 6.30 | 6.50 |
| 950 | 6.30 | 6.50 |

#### 4.4.7 Total Solid Fat content

Total Solid fat content of the sample was estimated to be 1.83 %

#### 4.4.8 Specific Gravity of Sample

Weight of raw milk sample and treated milk sample was found to be same that is 1.0325.

## 5 CONCLUSION

TiO_2_ thin films doped with zinc and copper were synthesized using sol-gel process and characterized using FTIR and SEM. FTIR showed the presence of Zn, Ag, and TiO_2_. While SEM showed a pore size of 0.5μm on the surface. Treatment of water with TiO_2_ thin films doped with zinc and copper increased the time required for methylene blue reduction, decreased the bacterial count by five log folds. Treatment of milk sample with TiO_2_ thin films doped with zinc and zinc with silver also increased the time required for methylene blue reduction. Treated milk didn’t curdle when boiled and also acidity wasn’t very high. The bacterial count had reduced to two digits. Tests for detection of cane sugar, hydrogen peroxide, starch, cellulose, sodium chloride showed no colour change indicating that TiO_2_ thin films don’t affect quality of milk. pH of sample was stable over period of time that is 6.61. Lastly total solid fat content of the sample was estimated to be 1.83 % and weight of raw milk sample and treated milk sample was found to be same that is 1.0325.

These results show the potential of using TiO_2_ thin films in reducing the microbial load and extending shelf life of milk.

Further these experiments can be carried out with other combination of metals as either dopants or base metals and also profile of various species of microorganisms can be studied in detail. In detail toxicology analysis of milk can be done by using invitro studies. This experiment was done on lab scale using glass microslide but on large scale it becomes obsolete as coating needs to be done large surface area which could be costly. So options of coating on glass rod and dipping in milk can and leaving it in the can from the time milk is drawn till its processed is one of the options and other option could be coating entire inner surface of milking vessel with film material. Both has its set of problems, in first case it needs to be addressed whether any metal residue is left in milk and in second case it needs to be seen if coating is reactive with material of milking vessel or any residue is left in this case also. If parameters are fine, then it is safe for human consumption.

